# CAT, AGTR2, L-SIGN and DC-SIGN are potential receptors for the entry of SARS-CoV-2 into human cells

**DOI:** 10.1101/2021.07.07.451411

**Authors:** Dongjie Guo, Ruifang Guo, Zhaoyang Li, Yuyang Zhang, Wei Zheng, Xiaoqiang Huang, Tursunjan Aziz, Yang Zhang, Lijun Liu

**Author notes:** Corresponding author. College of Life and Health Sciences, Northeastern University, Shenyang, 110169, China., *E-mail address:* (L. Liu). These authors contributed equally to this work (Dongjie Guo, Ruifang Guo).

## Abstract

Since December 2019, the COVID-19 caused by SARS-CoV-2 has been widely spread all over the world. It is reported that SARS-CoV-2 infection affects a series of human tissues, including lung, gastrointestinal tract, kidney, etc. ACE2 has been identified as the primary receptor of the SARS-CoV-2 Spike (S) protein. The relatively low expression level of this known receptor in the lungs, which is the predominantly infected organ in COVID-19, indicates that there may be some other co-receptors or alternative receptors of SARS-CoV-2 to work in coordination with ACE2. Here, we identified twenty-one candidate receptors of SARS-CoV-2, including ACE2-interactor proteins and SARS-CoV receptors. Then we investigated the protein expression levels of these twenty-one candidate receptors in different human tissues and found that five of which CAT, MME, L-SIGN, DC-SIGN, and AGTR2 were specifically expressed in SARS-CoV-2 affected tissues. Next, we performed molecular simulations of the above five candidate receptors with SARS-CoV-2 S protein, and found that the binding affinities of CAT, AGTR2, L-SIGN and DC-SIGN to S protein were even higher than ACE2. Interestingly, we also observed that CAT and AGTR2 bound to S protein in different regions with ACE2 conformationally, suggesting that these two proteins are likely capable of the co-receptors of ACE2. Conclusively, we considered that CAT, AGTR2, L-SIGN and DC-SIGN were the potential receptors of SARS-CoV-2. Moreover, AGTR2 and DC-SIGN tend to be highly expressed in the lungs of smokers, which is consistent with clinical phenomena of COVID-19, and further confirmed our conclusion. Besides, we also predicted the binding hot spots for these putative protein-protein interactions, which would help develop drugs against SARS-CoV-2.

## 1. Introduction

COVID-19 is an ongoing pandemic caused by the severe acute respiratory syndrome coronavirus 2 (SARS-CoV-2) [1]. As of June 30^th^, 2021, more than 180 million confirmed cases and 3,934,252 deaths worldwide have been reported, and the number of infected individuals is still increasing. Until now, the cell entry mechanism of SARS-CoV-2 has not been fully elucidated, and no effective drugs has been found for treating COVID-19. Therefore, intensive researches are urgently needed to elucidate the mechanisms of SARS-CoV-2 entry into the host, thereby providing a promising target for developing specific antiviral drugs.

It is reported that the cell entry of SARS-CoV-2 depends on the binding of the spike (S) protein to the primary functional receptor ACE2 and S protein cleaving and priming by host cell proteases TMPRSS2 [2–4]. However, the above mechanism could not fully explicate the clinical phenomena of SARS-CoV-2 infection. According to large-scale clinical studies, the main infected organ of COVID-19 is lung, in which the expression of ACE2 is relatively low [5–8]. And only 11.4%, 3.2%, and 8.64% patients have gastrointestinal tract, kidney and testis dysfunction respectively, while these tissues with extremely abundant expression of ACE2 [9–11]. In addition, TMPRSS2 also has a relatively low expression level in lung, colon, and kidney [12]. It remains unclear why it is the lung, rather than other tissues with higher ACE2 expression is mainly infected. Notably, a recent study reported that the administration of ACE2 inhibitors showed no association with clinical outcomes among COVID-19 patients [13]. Based on the above, there may be other factors assisting in virus entry. We hypothesized that some ACE2-interacting proteins which were specifically expressed in lung and other infected tissues and also had high binding capacities with S protein might be as the co-receptors of ACE2 to synergistically bind to S protein and further promote SARS-CoV-2 entry into host cells. Recently, neuropilin-1 is identified as a co-factor involved in ACE2-mediated SARS-CoV-2 infection [14, 15]. On the other side, besides the co-receptor of ACE2, we also intended to search other novel receptors. SARS-CoV-2 and SARS-CoV share a high sequence identity between their S proteins, and their cell entry is both dependent on ACE2 and TMPRSS2 [16, 17], which indicates that the other receptors of SARS-CoV are also likely to be the SARS-CoV-2 receptors.

In this study, we reported that four potential receptors, of which two ACE2-interacting proteins CAT, AGTR2, and two SARS-CoV receptors L-SIGN, DC-SIGN not only had specifically expression levels and in lung and other affected organs in COVID-19, but also had higher binding affinities with S protein than ACE2. Additionally, we also observed that the expression levels of AGTR2 and DC-SIGN were associated with smoking status. Given these observations, we suggested that CAT, AGTR2, L-SIGN and DC-SIGN could be the co-receptors or alternative receptors of SARS-CoV-2, and the binding hot spots we predicted at the receptors-S protein binding interfaces probably be the potential targets for treatment of COVID-19.

## 2. Methods

### 2.1. Online resources

Two databases, BioGRID [18] and STRING [19] were searched to identify potential ACE2-interacting proteins. The data of normal lung tissues in microarray datasets GSE10072, GSE123352, and GSE32863 from Gene Expression Omnibus (GEO) were analyzed for exploring the correlations between the expression levels of potential receptors and smoking status. After excluding the influence of variables such as age and race, a total of 138 samples were analyzed in smoking status, of which 30 were from GSE10072 (including 15 smokers and 15 non-smokers), 94 were from GSE123352 (including 47 smokers and 47 non-smokers), and 18 were from GSE32863 (including 9 smokers and 9 non-smokers). The protein and mRNA expression data of potential receptors were obtained from the Human Protein Atlas (HPA) portal (https://www.proteinatlas.org/). The experimental structure of AGTR2 was downloaded from the Protein Data Bank (PDB, http://www.rcsb.org/). The subcellular locations of potential receptors were acquired from UniProt (https://www.uniprot.org/). The amino acid sequences of all proteins were obtained from Ensembl (https://asia.ensembl.org/index.html). Data management using R 4.0.0, statistical analysis and data visualization using GraphPad Prism 8.0.2.

### 2.2. Protein structures prediction

The predicted structure of S protein was downloaded from the zhanglab (https://zhanglab.ccmb.med.umich.edu/COVID-19/) and only the structure part which belongs to residues 1-685 of S1 was retained for further analysis [17]. The structures of potential receptors other than AGTR2 were predicted by I-TASSER [20] (Table S1). For each protein, five models were generated and the model with the highest C-score was selected as the best one and used for the following analysis.

### 2.3. Protein-protein docking, binding affinity calculation, and hot spots prediction

Protein-protein docking between SARS-CoV-2 S protein and the putative receptors was performed by HADDOCK, and PRODIGY was used to predict the binding affinity for the best complexes formed in each docking experiment [21, 22]. Based on the complex models, the KFC2 server was used to predict binding hot spots within receptors-S interfaces by recognizing structural features indicative of important binding contacts [23]. Data visualization was accomplished using PyMOL.

### 2.4. Statistical analysis

Student’s t-test was used to compare mRNA expression of potential receptors between non-smoking and smoking samples. For all analyses, data were plotted as mean ± SEM, and a *P*-value < 0.05 was regarded as statistically significant.

## 3. Results

### 3.1. Identification of the candidate receptors for SARS-CoV-2

In order to find candidate receptors of SARS-CoV-2 from two clues, ACE2-interacting proteins and SARS-CoV receptors, we searched for protein-protein interaction databases. Firstly, In BioGRID and STRING [18, 19], thirty proteins were shown to interact with ACE2 (Figure. 1). Among these proteins, except that AGT was from both databases, TEX101, CALM1, PDZK1, VIM, SNX17, TMPRSS11D, TMPRSS11A, SHANK1, ISYNA1, DEFA5, SLC6A19, AAMP, MDM2, CIT, HSPA5, CDSN, CAT, TMPRSS2, TFRC, and HRAS were from BioGRID (Fig. 1A), while AGTR1, AGTR2, REN, MME, Mep1A, Mep1B, PRCP, XPNPEP2, and DPP4 were from STRING (Fig. 1B). Moreover, L-SIGN, DC-SIGN, and CD13 were previously reported to be receptors for SARS-CoV [24–26], and thus were selected as candidate receptors.

**Figure 1.**
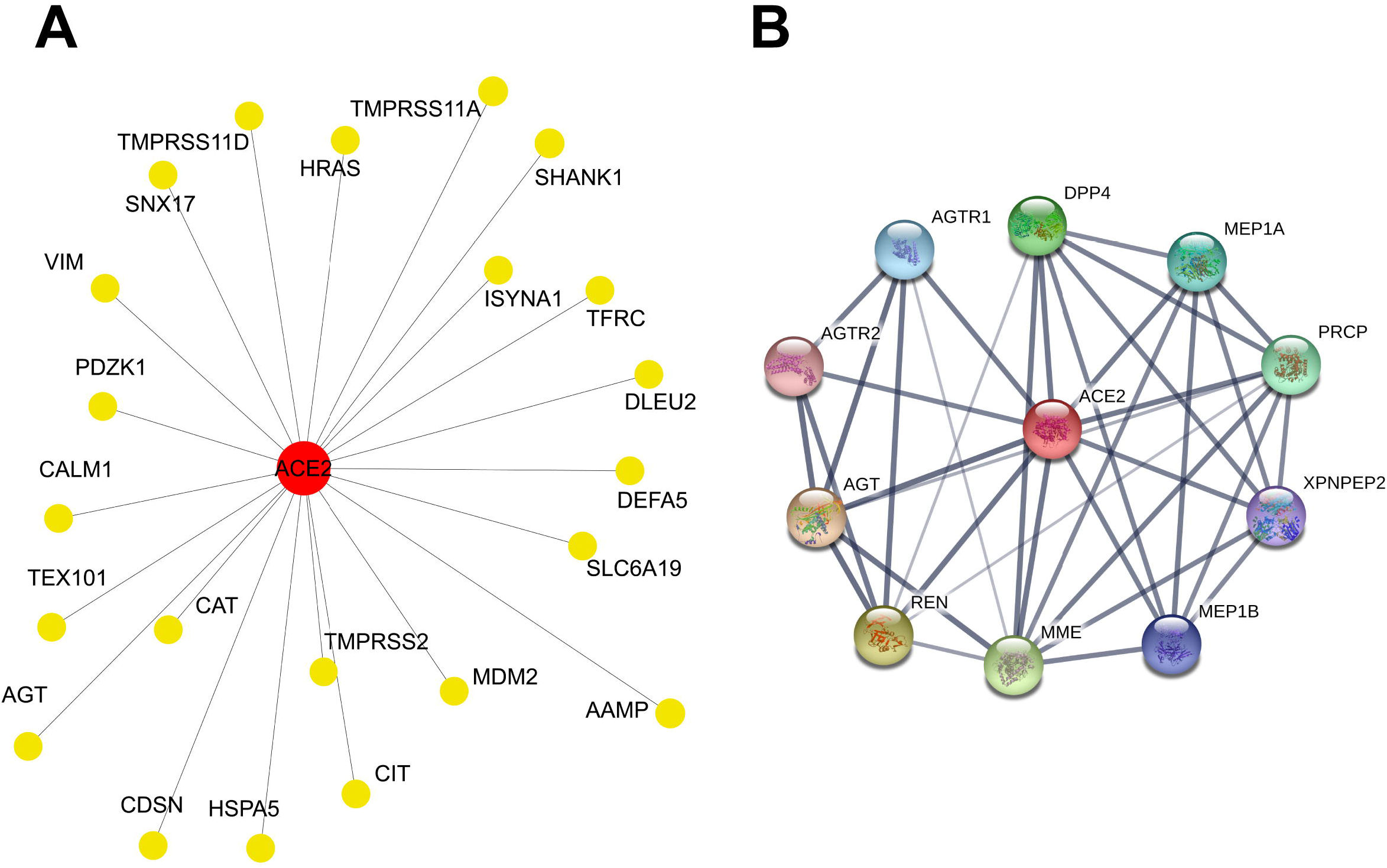
Protein interaction networks of ACE2. (A) Protein interaction networks of ACE2 from BioGRID. Among ACE2 interactors, DLEU2 is a long noncoding RNA, and left ones are proteins. (B) Protein interaction networks of ACE2 from STRING.

Therefore, a total of thirty-three candidate receptors that might bind to SARS-CoV-2 S protein were initially considered.

Considering that only membrane proteins on host cells could act as cell entry receptors of virus, we searched UniProt [27] to figure out the subcellular localization of these candidates. The results showed that CALM1, SNX17, TMPRSS11A, SHANK1, ISYNA1, DEFA5, CIT, REN, MEP1A, PRCP, MDM2, and AGT were not membrane proteins and hence were excluded from further analysis. Therefore, the remained twenty-one candidate receptors were evaluated.

### 3.2. The expression levels of CAT, MME, L-SIGN, DC-SIGN, and AGTR2 were specifically high in lung and other tissues affected in COVID-19

It is reported that SARS-CoV-2 infection affects a series of human tissues, including lung, gastrointestinal tract, kidney, testis, epididymis, cerebral cortex, gallbladder, etc [5, 9, 11, 28-32]. So we investigated the protein expression levels of the twenty-one candidate receptors selected above from HPA and found that five of which CAT, MME, L-SIGN, DC-SIGN, and AGTR2 were specifically expressed in these tissues, whereas the remaining candidates were not specifically expressed in affected tissues such as lung (Fig. 2 and Fig. S1). The results showed that CAT was moderately expressed in duodenum and kidney, followed by lung and small intestine (Fig. 2A). MME was highly expressed in small intestine, duodenum and kidney, moderately expressed in epididymis and gallbladder, followed by lung (Fig. 2B). L-SIGN could be detected in lung, colon and cerebral cortex, and DC-SIGN was moderately expressed in lung (Fig. 2C and D). Although there is no protein expression annotation, the mRNA level of AGTR2 was relatively high in lung, with high tissue specificity (Fig. 2E). Taken together, the expression levels of CAT, MME, L-SIGN, DC-SIGN and AGTR2 were consistent with the clinical symptoms in COVID-19, which indicate that these five proteins may be potential receptors for SARS-CoV-2.

**Figure 2.**
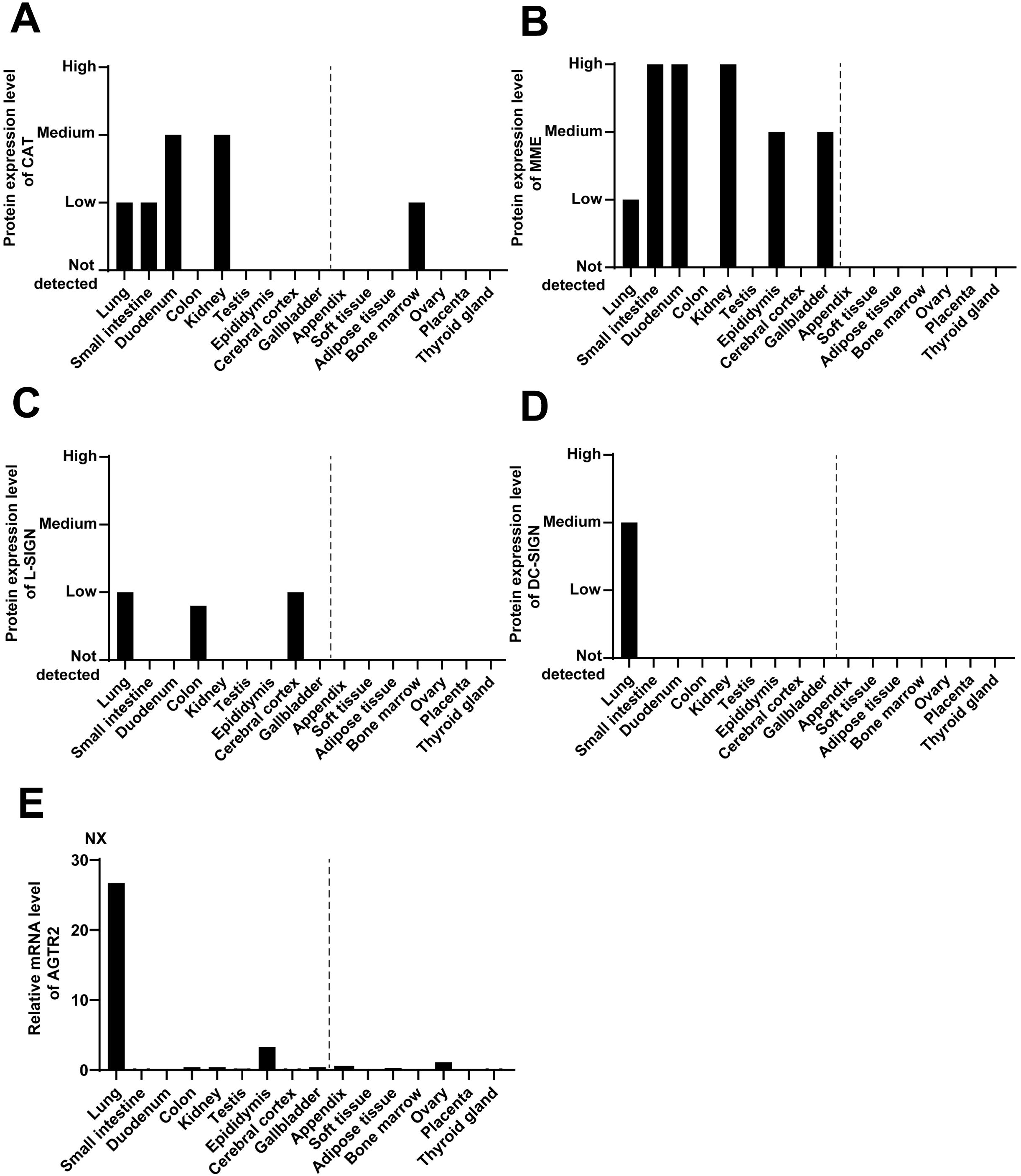
Expression levels of the potential receptors in different human tissues. (A-D) Bars show protein expression levels of CAT, MME, L-SIGN, and DC-SIGN in clinically affected and unaffected human tissues by SARS-CoV-2. (E) Bars show mRNA expression level of AGTR2 in clinically affected and unaffected human tissues by SARS-CoV-2. Data are from NCBI. Abbreviation: NX, Normalized expression. On the left of the dotted line are affected tissues, and on the right are unaffected tissues.

### 3.3. L-SIGN, CAT, AGTR2, and DC-SIGN showed higher binding affinities with SARS-CoV-2 S protein than ACE2

To quantify the interactions between the potential receptors and SARS-CoV-2 S protein, we modeled their interactions and predicted the binding affinities based on the complex models [21, 22, 33]. In coronavirus (CoVs), the entry process is mainly mediated by S protein [34]. The S protein would, in most cases, be cleaved by host proteases into the S1 and S2 subunits, which are responsible for receptor recognition and membrane fusion, respectively [35]. S1 can be further divided into an N-terminal domain (NTD) and a C-terminal domain (CTD). Although CTD is usually as receptor-binding domain (RBD) for CoVs, other CoVs, like mouse hepatitis CoV, engages the receptor with its NTD [36–38]. Consequently, we employed the S1 for further protein interaction simulations based on three-dimensional structures.

Given that the crystal structure of the receptor binding domain of SARS-CoV-2 in complex with ACE2 has been solved, we performed a simulation of 3D structure based protein-protein interaction only for candidate receptors (Fig. 3A and Fig. S2). The results showed that the binding free energy (ΔG) of ACE2 and RBD (S) complex was −12.4 kcal/mol. Strikingly, the binding affinities of L-SIGN, CAT, AGTR2 and DC-SIGN with S1 were higher than ACE2, with binding free energy (ΔG) of −17, −16.9, −15.8, and −15.5 kcal/mol, respectively (Fig. 3B). While the binding affinity of MME with S1 was lower than ACE2, with a binding free energy (ΔG) of −11.8 kcal/mol (Fig. 3B). Furthermore, we observed that CAT and AGTR2 bound to S1 in different regions with ACE2 conformationally, suggesting that these two proteins are likely capable of the co-receptors of ACE2 (Fig.3A). Based on the above analyses, we considered that L-SIGN, CAT, AGTR2 and DC-SIGN served as the potential receptors for SARS-CoV-2.

**Figure 3.**
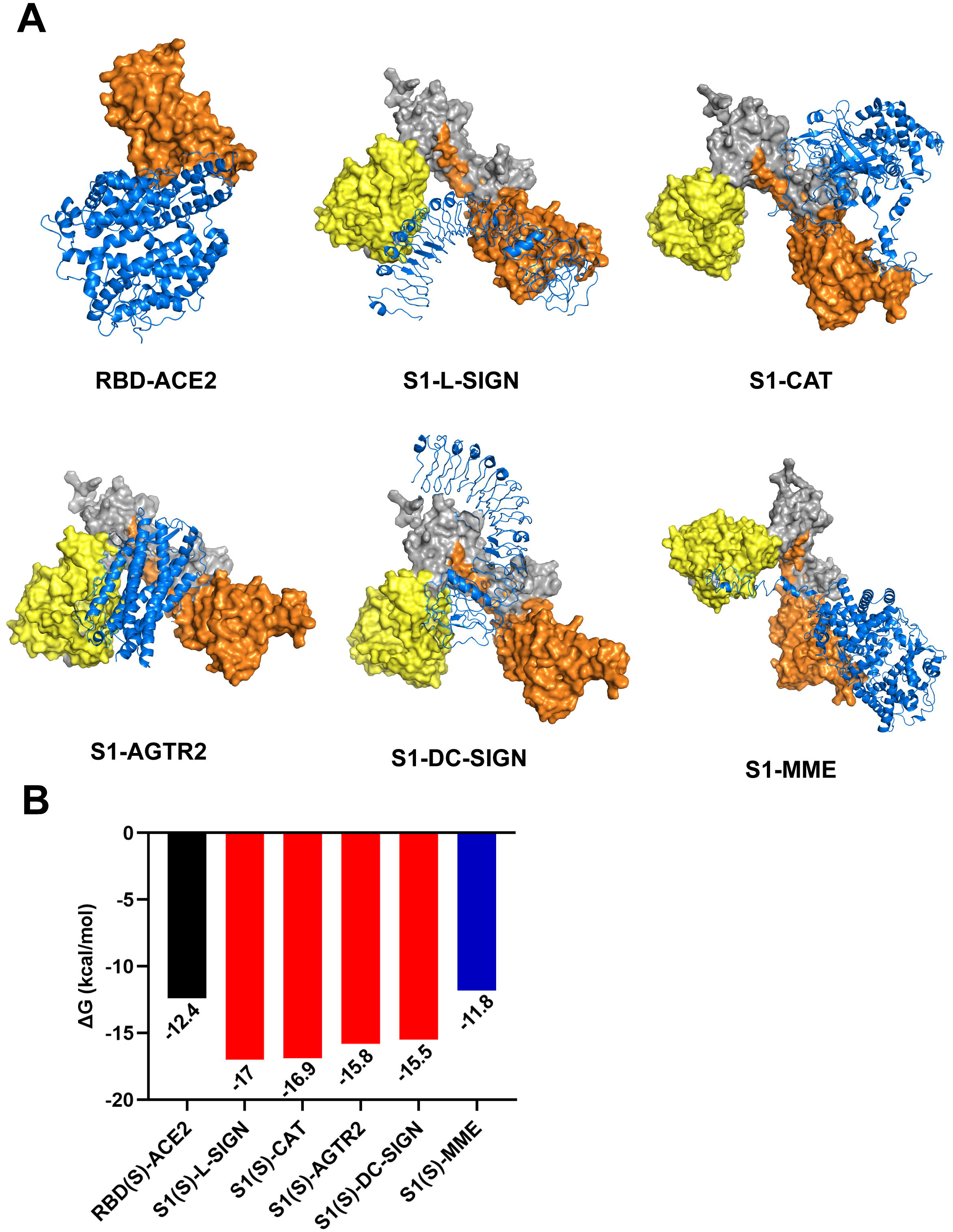
Molecular docking of potential receptors and SARS-CoV-2 S protein. (A) The 3D structure of RBD (orange) and ACE2 (blue) complex from PDB (ID: 6LZG). Predicted 3D structures of the complex formed by S1 (yellow, orange and gray) and potential receptors (blue) by HADDOCK. (B) The binding affinities (ΔG) of ACE2-RBD (S) complex (black column) and potential receptors-S protein complexes (red and blue columns).

### 3.4. The expression levels of AGTR2 and DC-SIGN were associated with smoking status

According to previous reports, the proportion of current smoker was statistically significant higher in lethal group compared to the non-lethal group [39], which indicating that the disease severity of COVID-19 was positively connected with smoking status. To explore whether the mRNA expression levels of these potential receptors correlate with smoking status, We analyzed the mRNA expression data in normal lung tissue from three microarray data sets, i.e. GSE10072, GSE32863 and GSE123352. The results showed that the mRNA expression level of AGTR2 in the smoking group was significantly higher than that in the non-smoking group in GSE10072 (*P* = 0.0150) and GSE32863 (*P* = 0.0387), a similar trend was observed in GSE123352 (*P* = 0.0979) (Fig. 4C). We also observed a higher expression level of DC-SIGN in smokers compared with non-smokers in both GSE10072 (*P* = 0.0295) and GSE32863 (*P* = 0.0215) (Fig. 4D). However, no significant difference in the expression levels of L-SIGN and CAT were observed between smoking and non-smoking groups (Fig. 4A and B). Collectively, the mRNA expression levels of AGTR2 and DC-SIGN in lung were associated with smoking status, which is consistent with clinical symptoms of COVID-19, and further confirmed our conclusion above.

**Figure 4.**
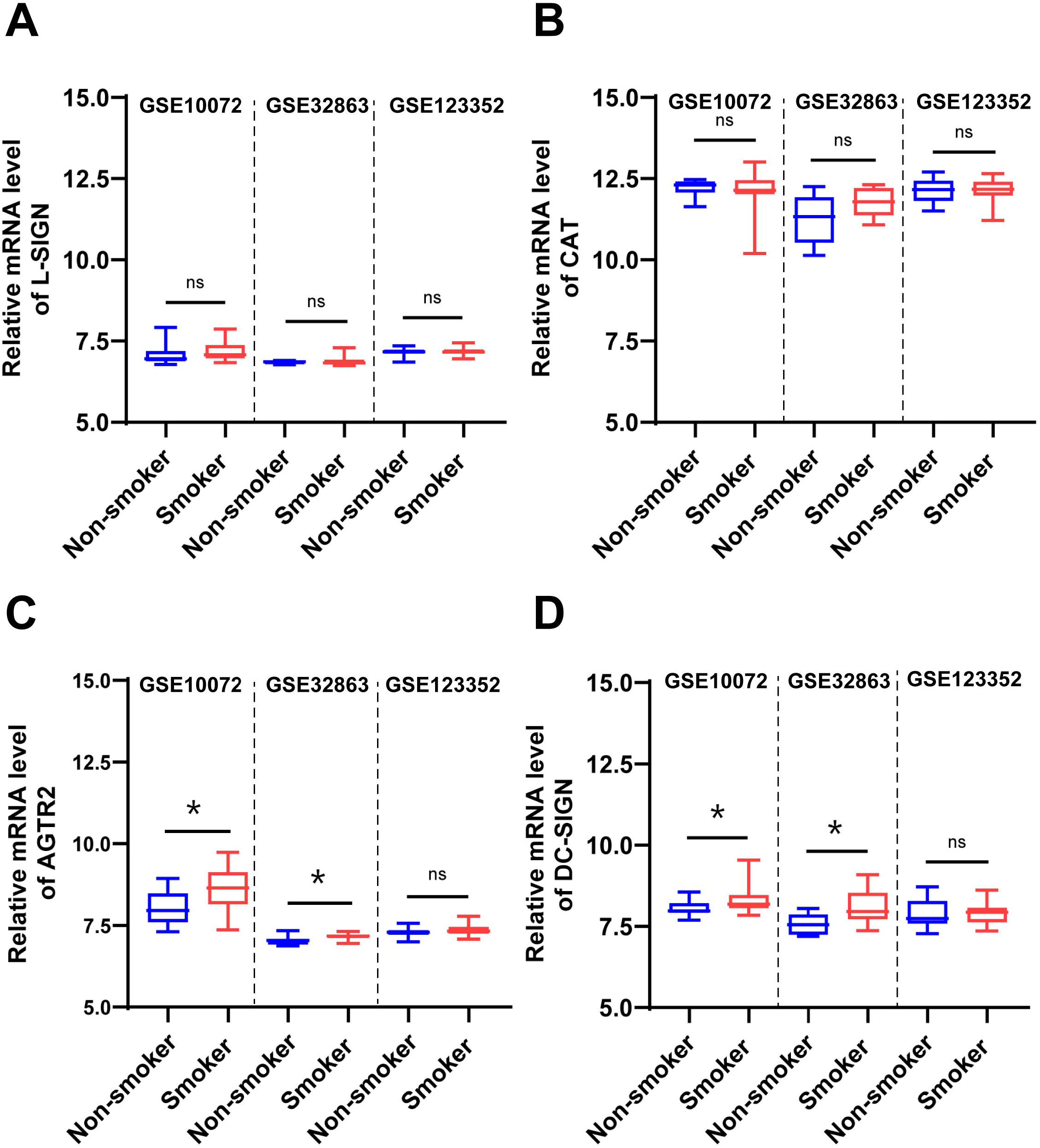
mRNA expression profiles of L-SIGN, CAT, AGTR2 and DC-SIGN in smoking and non-smoking lung. (A) mRNA expression level of L-SIGN (*P*=0.3678 in GSE10072; *P*=0.9895 in GSE32863; *P*=0.2283 in GSE123352), (B) CAT (*P*=0.8302 in GSE10072; *P*=0.1092 in GSE32863; *P*=0.9801 in GSE123352), (C) AGTR2 (*P* = 0.0150 in GSE10072; *P* =0.0387 in GSE32863; *P*=0.0979 in GSE123352), (D) DC-SIGN (*P*=0.0295 in GSE10072; *P*=0.0215 in GSE32863; *P* =0.6447 in GSE123352) in normal lung tissues of smoking and non-smoking group. Data is shown in mean ± SEM; GSE10072, n=15; GSE123352, n=47; GSE32863, n=9; \**P*<0.05, \*\**P*<0.01, \*\*\**P*<0.001 (Student’s t test).

### 3.5. Prediction of binding hot spots in the complex of potential receptors and S protein

When protein molecules form a complex, a subset of residues in the interface that contribute the most to binding is regarded as hot spots, which are crucial for preserving protein function, maintaining the stability of protein association, and developing antiviral drugs. Hence, in order to better understand the interactions between potential receptors and S protein and find novel targets, we used the KFC2 server to computationally predict the binding hot spots in potential receptors and S protein complex. Firstly, the binding hot spots on potential receptors identified as follows: 1) A31, I42, Q53, V55, V56, L355, F356, D360, R363 on CAT, 2) P75, I81, L164, P165, C169, L176, F179 on AGTR2, 3) S345, C381, D382, Y386, W387, I388, C389, K390 on L-SIGN, 4) R309, S338, F339, Y342, F359, G361, N362, I376, E389, F391, L392, S393 on DC-SIGN (Fig. 5B, C and Fig. 6A, B), may be potential targets for the treatment of COVID-19. Then further analysis was performed to compared the interface residues on S protein. We found that on S protein, the binding sites of CAT and AGTR2 were located in different regions with ACE2 (Fig. 5). Therefore, the two ACE2-interacting proteins, CAT and AGTR2, may synergistically bind to S protein as the co-receptors of ACE2. Taken together, these binding hot spots in potential receptors-S protein complex might be novel drug targets against SARS-CoV-2.

**Figure 5.**
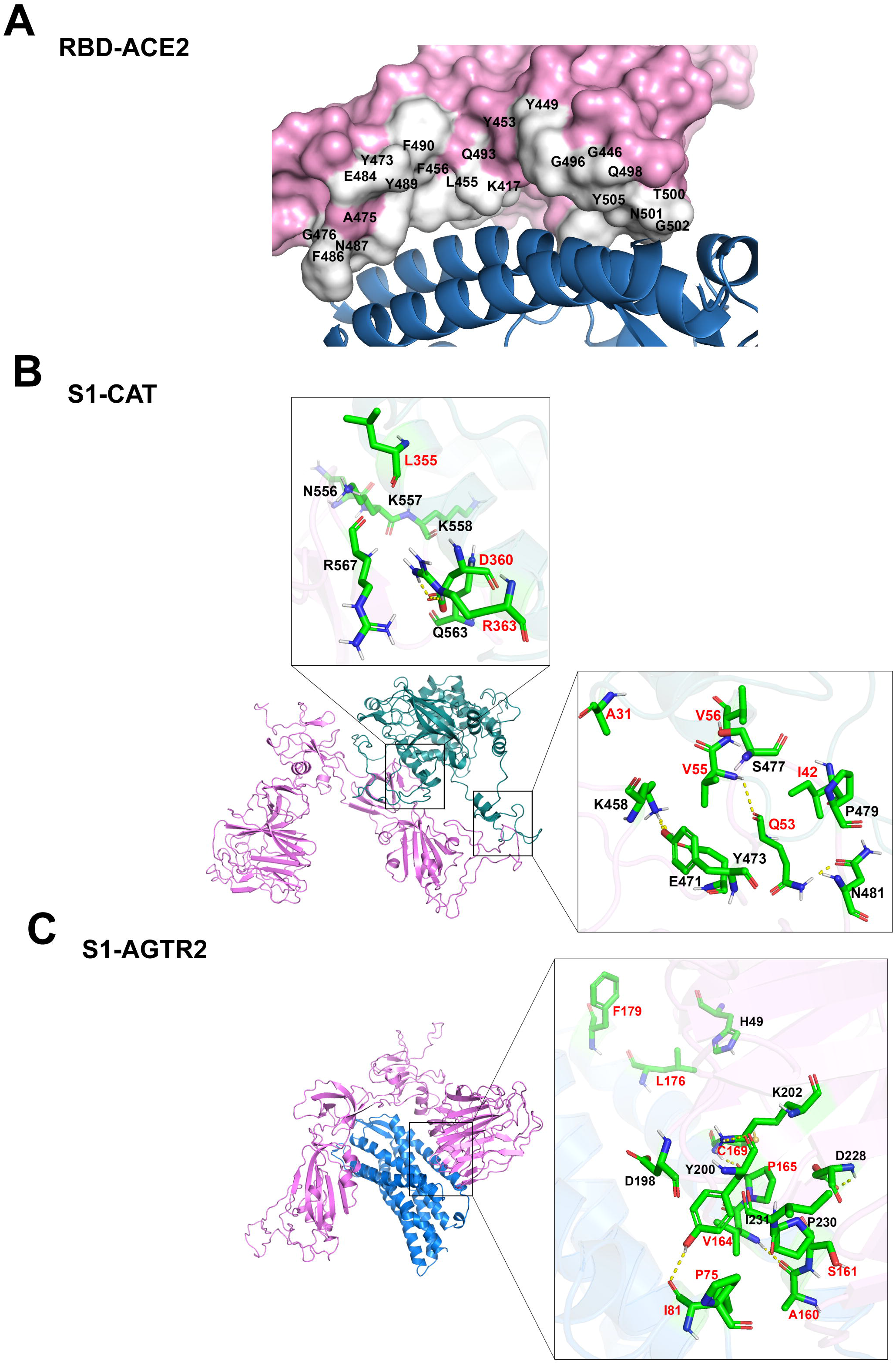
The binding hot spots in RBD-ACE2 and potential receptors-S1 complexes. (A) Residues on RBD that interact with ACE2 are marked. (B&C) In the complexes of CAT, and AGTR2 with S1, binding hot spots on CAT, AGTR2 are indicated in red fonts, and on S1 are indicated in black fonts.

**Figure 6.**
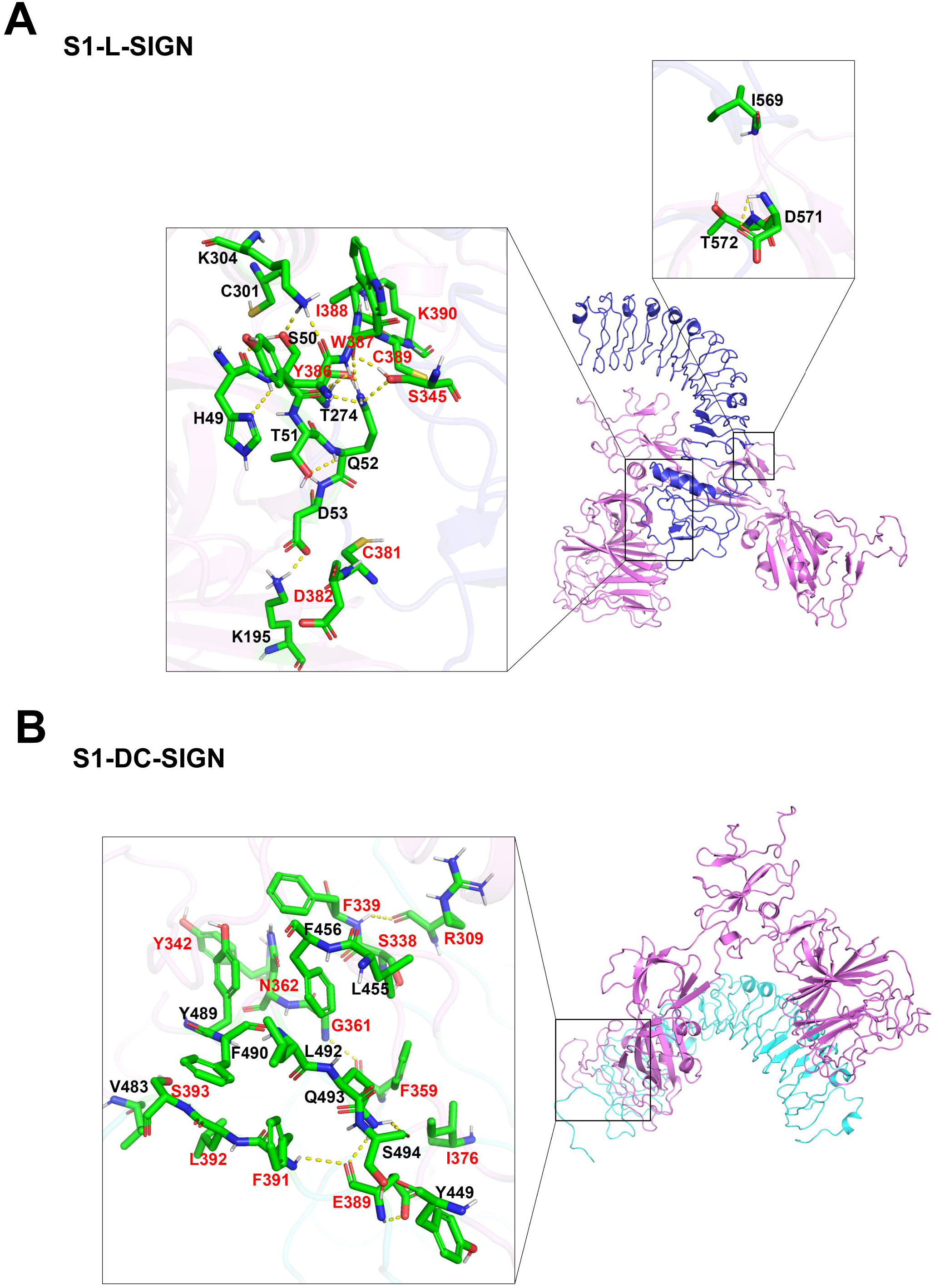
The binding hot spots in potential receptors-S1 complexes. (A&B) In the complexes of L-SIGN, and DC-SIGN with S1, binding hot spots on L-SIGN and DC-SIGN are indicated in red fonts, on S1 are indicated in black fonts.

## 4. Discussion

In COVID-19, the expression levels of the known receptors ACE2 and TMPRSS2 in affected tissues are not consistent with clinical symptoms, which suggests that there may be other receptors involved in the cellular entry of SARS-CoV-2. Our study demonstrates that host cell entry of SARS-CoV-2 possibly depends on two ACE2-interacting proteins CAT, AGTR2, and two SARS-CoV receptors L-SIGN and DC-SIGN. The results have crucial implications for our understanding of the molecular and cellular effects of SARS-CoV-2. Moreover, binding hot spots in these potential receptors and S protein complex reveal targets for therapeutic interventions.

CAT, a key antioxidant enzyme [40], is specifically expressed in lung, kidney, intestine and other affected tissues in COVID-19. Overexpression of CAT has been reported to reduce renal oxidative stress, prevent hypertension and show a correlation with ACE2 expression [41]. Intriguingly, ACE2 deficiency can increase NADPH-mediated oxidative stress in kidney, suggesting a possible link with CAT [42]. Furthermore, protein-protein interaction simulations revealed that CAT could bind to S protein with a higher binding affinity than ACE2. The another ACE2-interacting protein, AGTR2, belongs to the G protein coupled receptor 1 family, and functions as a receptor for angiotensin II. It is highly expressed in lung with a high tissue specificity. Previous studies have shown that ACE2 and the AGTR2 protect mice from severe acute lung injury induced by acid aspiration or sepsis [43], which illustrates their functional similarities. In particular, AGTR2 could bind to S protein with a higher binding affinity than ACE2 and different binding sites from ACE2. Conformably, a recent study also mentioned that AGTR2 had a higher binding affinity with SARS-CoV-2 S protein than ACE2 [44]. Moreover, our results also suggested that the mRNA expression level of AGTR2 was associated with smoking status, which consistent with the clinical phenomenon. Thus, we considered that CAT and AGTR2 act as the coreceptors of ACE2 for stabilizing the binding of ACE2 and S protein and promoting the cells entry of SARS-CoV-2. Consistently, a recent study found that kidney injury molecule-1 (KIM1) and ACE2 bind in different regions of SARS-CoV-2 through molecular simulations, indicating that they have the potential to synergistically promote virus invasion [45].

Previous studies demonstrated that L-SIGN and DC-SIGN could interact with SARS-CoV S protein and mediate virus infection [46]. Now our proteinprotein docking results showed that L-SIGN and DC-SIGN could also bind to SARS-CoV-2 S protein, which is highly similar with SARS-CoV S protein. Moreover, loss of L-SIGN in mice significantly reduced SARS-CoV infection, further emphasizing the critical role of L-SIGN in SARS-CoV infection [47]. Furthermore, DC-SIGN and L-SIGN both appear to have higher affinities with SARS-CoV-2 S protein even than ACE2 (binding free energy (ΔG) −17 vs - 12.4 kcal/mol, −15.5 vs −12.4 kcal/mol respectively), suggesting that these two proteins may function as the alternative receptors independent of ACE2. Interestingly, unlike ACE2, which is present at relatively low levels in lung and other organs, DC-SIGN is highly expressed in lung. While L-SIGN is broadly expressed in lung, colon and cerebral cortex, indicating that it may function in multiple tissues as a broad-spectrum receptor. In addition, the mRNA expression level of DC-SIGN was related to smoking status. Consistently, previous studies have also shown that the expression level of DC-SIGN is higher in smokers than non-smokers [48]. These evidences further suggest that two SARS-CoV receptors, L-SIGN and DC-SIGN, may also play roles in the entry of SARS-CoV-2 into human cells. Based on the modeled potential receptors-S protein complex, we also predicted the binding hot spots, which would be helpful to repurpose or design drugs targeting these potential receptors. Currently, some inhibitors targeting these potential receptors can be found in DrugBank, such as CAT inhibitor Fomepizole, AGTR2 antagonist Tasosartan. In addition, L-SIGN inhibitor Dextran and DC-SIGN inhibitors quinoxalinones were also identified as potential drugs [49, 50]. Finally, our findings suggest that CAT, AGTR2, L-SIGN and DC-SIGN could be novel potential receptors for the entry of SARS-CoV-2 into human cells and the identified agents should be carefully considered in anti-SARS-CoV-2 usage.

## Acknowledgments

We thank Xiaoting Li (Columbia University, USA) for helpful advice on modeling analysis. We thank Dr. Qi Zhao (Northeastern University, China) for technical assistance on molecular docking. This study was financially supported by Northeastern University, PRC (N2020006), and Talent Project of Revitalizing Liaoning (XLYC1907052). The funders had no role in study design, data collection, analysis, decision to publish, or preparation of the manuscript.

## Supplemental information

Supplemental information includes two figures and one table.

## Author contributions

L.L. conceived the study, acquired funding, designed computational experiments, and assumed supervision. D.G. and R.G. drafted the manuscript. Y.Y.Z. analyzed the clinical symptoms of COVID-19. D.G., R.G., Z.L. and Y.Y.Z. identified the potential protein receptors. W.Z., X.H. and Y.Z. contributed to protein structure prediction. D.G. and Z.L. helped in protein expression analysis and protein-protein docking. D.G. and R.G. analyzed the correlation between receptor expression and smoking status. D.G., Z.L., and R.G. performed hot spot analysis. D.G. and R.G. created figures. D.G., R.G., T.A., Z.L., and L.L. edited the manuscript.

## Data availability

The data that support the findings of this study are available from the corresponding author upon reasonable request.

---

**Figure S1. Expression levels of the candidate receptors in different human tissues.**

Bars show protein expression levels of PDZK1, TEX101, VIM, TMPRSS11D, SLC6A19, AAMP, HSPA5, CDSN, TFRC, HRAS, TMPRSS2, AGTR1, MEP1B, XPNPEP2, DPP4, and CD13 in clinically affected and unaffected human tissues by SARS-CoV-2. Those on the left of the dotted line are affected tissues, those on the right are unaffected tissues.

**Figure S2. Predicted structures of S1 and candidate receptors.**

Structure of S1 was downloaded from the zhanglab (https://zhanglab.ccmb.med.umich.edu/COVID-19/). Structure of AGTR2 was downloaded from PDB (ID: 5UNG). And structures of CAT, MME, L-SIGN, DC-SIGN were predicted by C-I-TASSER. TM-score and C-score of each structure model was listed below.

**Table S1. Parameters resulting in the 3D structure of the candidate receptors predicted by I-TASSER.**

