## Supplementary figures and images for "CAT, AGTR2, L-SIGN and DC-SIGN are potential receptors for the entry of SARS-CoV-2 into human cells"

### Supplemental Figure 1

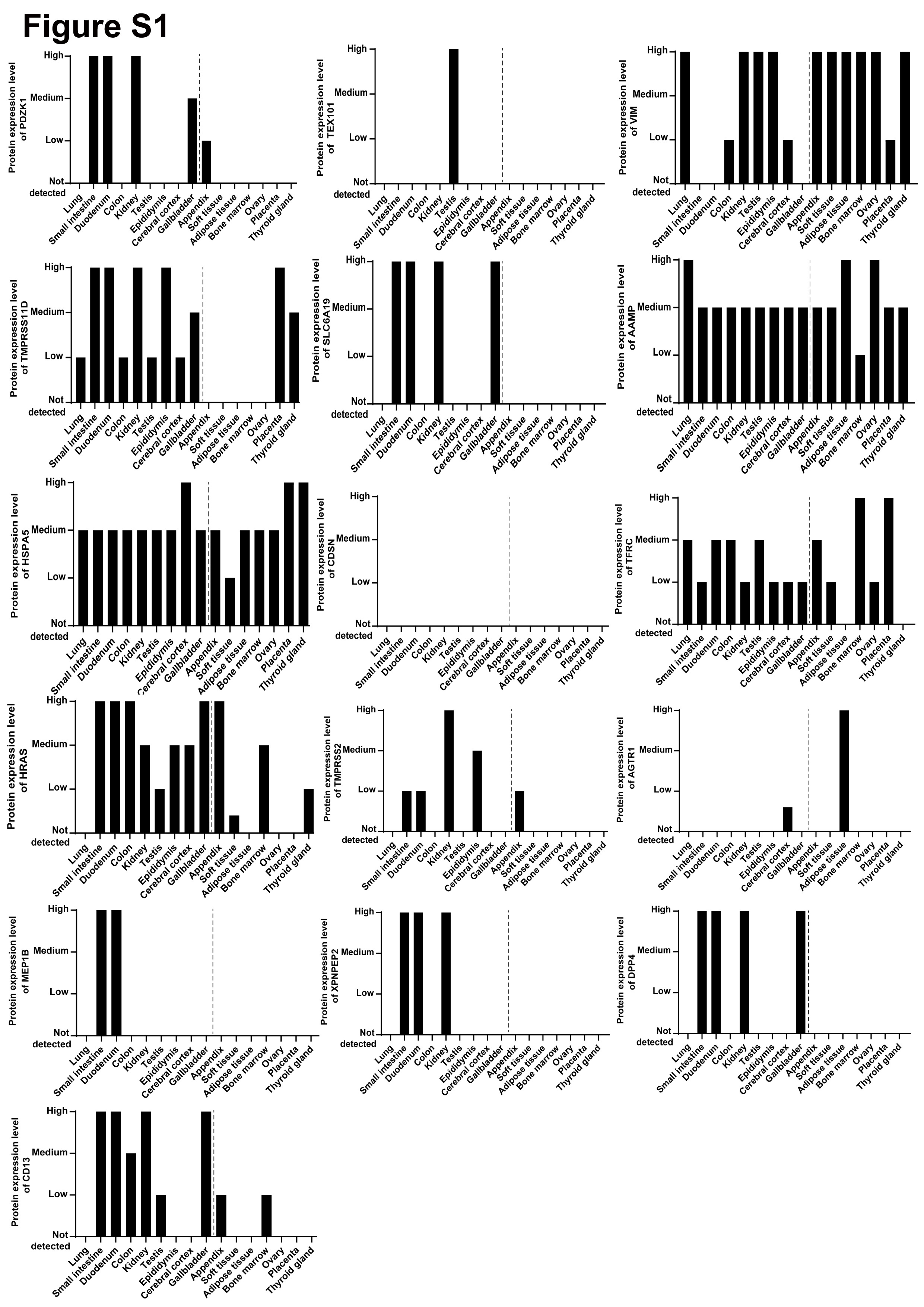

### Supplemental Figure 2

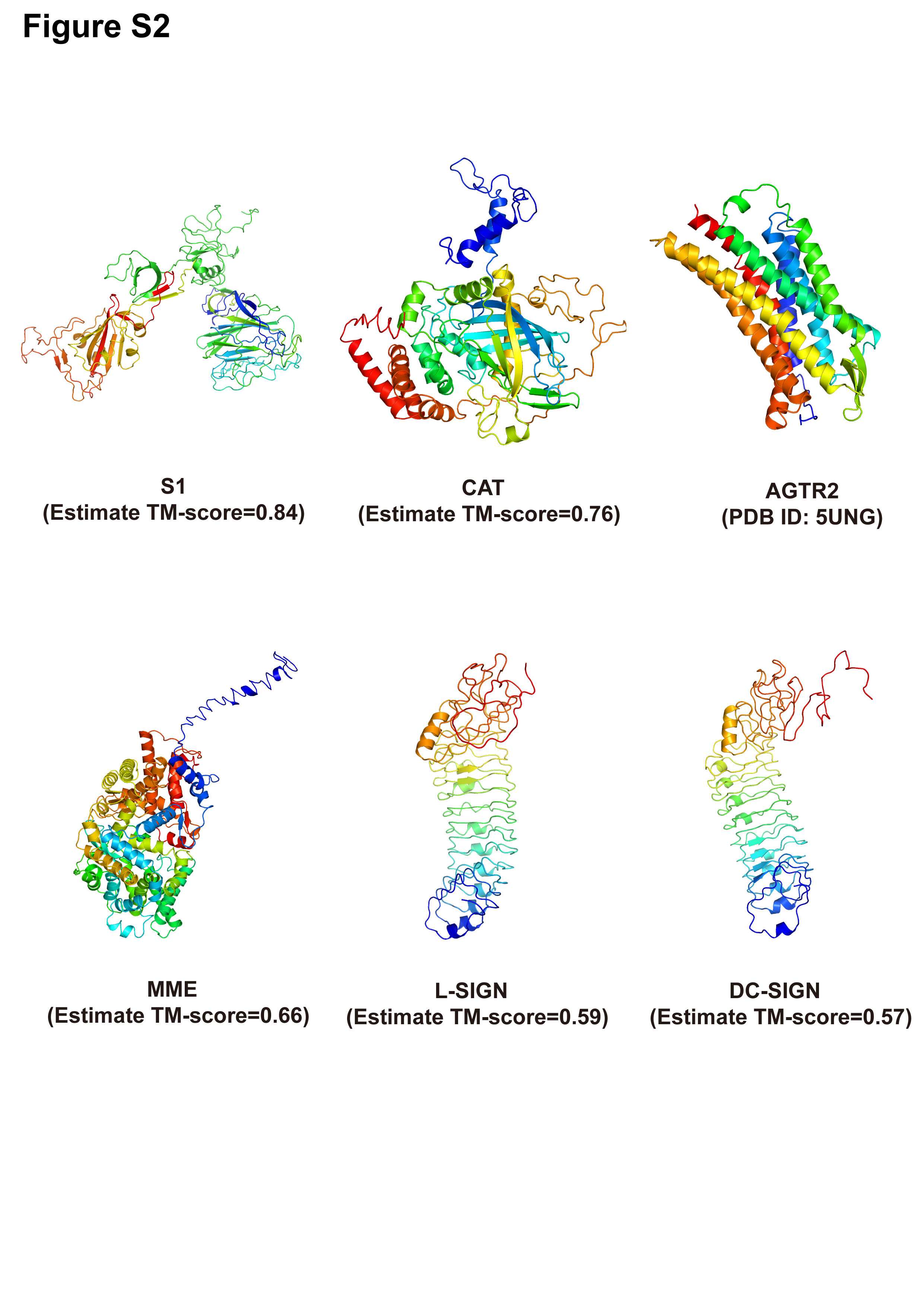
