## Supplemental Table 1 for "CAT, AGTR2, L-SIGN and DC-SIGN are potential receptors for the entry of SARS-CoV-2 into human cells"

**Table S1.** **Parameters resulting in the 3D structure of the candidate receptors predicted by I-TASSER**

| **Candidate**  **receptors** | **Protein ID** | **C-score** | **Estimate TM-score** |
| --- | --- | --- | --- |
| CAT | ENSP00000241052 | 0.34 | 0.76 |
| MME | ENSP00000418525 | -0.38 | 0.66 |
| L-SIGN | ENSP00000316228 | -0.98 | 0.59 |
| DC-SIGN | ENSP00000315477 | -1.16 | 0.57 |
